# Impact of clonal architecture on clinical course and prognosis in patients with myeloproliferative neoplasms

**DOI:** 10.1101/2022.09.24.509318

**Authors:** Damien Luque Paz, Michael S. Bader, Ronny Nienhold, Shivam Rai, Tiago Almeida Fonseca, Jan Stetka, Hui Hao-Shen, Gabi Mild-Schneider, Jakob R. Passweg, Radek C. Skoda

## Abstract

Myeloproliferative neoplasms (MPNs) are caused by a somatic gain-of-function mutation in one of three “disease driver” genes *JAK2, MPL* or *CALR*. About half of MPN patients also carry additional somatic mutations that modify the clinical course. The order of acquisition of these gene mutations has been proposed to influence the phenotype and evolution of the disease. We studied 50 *JAK2*-V617F-positive MPN patients who carried at least one additional somatic mutation and determined the clonal architecture of their hematopoiesis by sequencing DNA from single cell derived colonies. In 22 of these patients we also side-by-side applied Tapestri single-cell DNA sequencing (scDNAseq) with cells from the same blood sample. The clonal architectures derived by the two methods showed good overall concordance. scDNAseq showed higher sensitivity for mutations with low variant allele fraction, but had more difficulties distinguishing between heterozygous and homozygous mutations. By unsupervised analysis of clonal architecture data from all 50 MPN patients we defined 4 distinct clusters that differed by the order of acquisition of the mutations, and the complexity of the subclonal structure. Cluster 4, characterized by more complex subclonal structure without a preferred order of acquisition, correlated with reduced overall survival, and in multivariate analysis represented a risk factor independent of the MPN subtype or the age at diagnosis. Our results suggest that deciphering the clonal architecture in patients with MPN that carry multiple gene mutations can improve the molecular prognostic stratification that until now was primarily based on the number and type of gene mutations.

## Introduction

*BCR-ABL1* negative Myeloproliferative neoplasms (MPNs) are clonal disorders of hematopoietic stem cells driven by a somatic gain-of-function mutation in either *JAK2, MPL* or *CALR*.^1–7^ MPN patients at diagnosis are assigned to one of three subtypes: polycythemia vera (PV), essential thrombocythemia (ET) and primary myelofibrosis (PMF). The prognosis of MPNs is mostly dependent on the presence or absence of complications such as thrombotic or hemorrhagic events, and on the progression to secondary myelofibrosis or acute myeloid leukemia.^8^ Next-generation sequencing (NGS) allowed efficiently detecting additional gene mutations, which are often associated with less favorable prognosis.^9,10^ Additional somatic mutations can be acquired before or after the driver gene mutation or can represent a separate clone,^11^ and the order of acquisition has been proposed to influence the phenotype and the evolution of the disease.^12^

Several methods have been developed to determine the subclonal structure in patients with more than one somatic mutation. Genotyping of colonies derived from single cells, grown either in semi-solid media (e.g. methylcellulose) or in liquid culture, is considered the gold standard for determining clonal architecture, but this approach is labor-intensive and difficult to use for high throughput applications for larger cohorts of patients. Methods for genotyping of single cells without expansion in culture have first been developed at the RNA level and more recently also on DNA level. The Tapestri single-cell DNA sequencing (scDNAseq) method was previously used to determine clonal architecture in patients with AML,^13,14^ but no direct comparison with genotyping of single cell colonies was performed. Both the colony based and single cell-based methods require that gene mutations have been previously identified, e.g. by targeted or whole exome/genome NGS.

In the present study, we compared the results derived by genotyping of single cell derived colonies side by side with the Tapestri scDNAseq method in MPN patients with *JAK2*-V617F and at least one additional somatic mutation. We used the clonal architecture data to assign the patients to one of 4 distinct clusters and analyzed their associations with clinical features and prognosis.

## Methods

### Patients and samples

Collection of blood samples and clinical data of MPN patients at the University Hospital Basel, Switzerland, was approved by the local Ethics Committee (Ethik Kommission Beider Basel) and written informed consent was obtained from all patients in accordance with the Declaration of Helsinki. The diagnosis of MPN was established according to the revised criteria of the World Health Organization.^21^

### Next-generation sequencing

The mutational profiles of patients was established by a targeted NGS panel of granulocyte DNA covering all exons of 68 genes as previously described.^9^ Patients with *JAK2*-V617F and one or more additional somatic mutations were selected for the study.

### Single-cell DNA sequencing

A vial of frozen peripheral blood mononuclear cells (PBMCs) was thawed and then split for scDNAseq and single cell liquid culture for genotyping of colonies. Tapestri scDNAseq (Mission Bio^™^) was performed with a custom panel of 97 amplicons covering all 31 mutated gene regions previously determined in the 22 selected patients by NGS,{Lundberg, 2014 #7024} and 66 amplicons covering classical mutational hotspots within the selected genes and in 2 additional genes (*CALR* and *RUNX1*) not mutated in any of the 22 patients. Briefly, libraries were built from 120’000 peripheral blood mononuclear cells (PBMCs) loaded into a microfluidic cartridge for a theorical final number of 10’000 cells captured, and quality was accessed by migration on a Bioanalyser DNA1000 chip (Agilent^®^). Finally, libraries were pooled for a 2×150 bp sequencing on an Illumina NovaSeq for a theorical number of 52 million reads per sample. Data were processed by the Tapestri Insight v2.2 software and a custom R script. Briefly, variant calling was performed with GATKv3.7 in each cell and variants were filtered in order to remove low confidence mutations. Uncertainties concerning homozygous versus heterozygous mutations due to potential allele drop-out were manually curated. Preparation of samples and libraries and bioinformatic analyses are detailed in Supplemental Methods.

### Genotyping of colonies

CD34-positive and Lin-negative cells from PBMC were single-cell sorted into 96-well plates containing 100 μL of free medium consisting of StemSpam SFEMII (Stemcell Technologies) with human cytokines SCF, IL-3, IL-6, IL-9, IL-11, TPO, G-CSF (20ng/mL each), GM-CSF (50ng/mL) and EPO (3U/mL) (Peprotech, Rocky Hill, NJ). Fresh media (50μL) was added at day 8 of culture. On day 14, DNA was extracted from colonies using Chelex 100 Resin (Cat. No. 143-2832, Bio-Rad Laboratories, Hercules, CA, USA), and known somatic mutations were genotyped either by allele specific PCR for *JAK2*-V617F or by Sanger sequencing of amplicons covering the mutated gene regions (details in Supplemental Methods).

### *JAK2*-V617F *quantification by digital PCR*

Aliquots of the cells used for scDNAseq library (unfractionated PBMCs) and for liquid culture (CD34-positive cells from PBMCs) were analyzed by digital PCR to determine the VAF for *JAK2*-V617F. Briefly, DNA was extracted using QIAmp DNA micro kit (Qiagen) and then loaded on a chip with 20’000 partitions for allele specific PCR using probes with FAM marker for wild type *JAK2* and VIC for *JAK2*-V617F (Quantstudio3D, ThermoFisher).

### Statistics

Accuracy of clonal architecture methodology to derive the *JAK2*-V617F VAF was accessed by a non-parametric Passing-Bablok regression with the digital PCR measurement as a reference. For unsupervised clustering, Factor Analysis of Mixed Data (FAMD) was performed using 10 clonal architecture parameters (see Supplemental Methods) and followed by a clustering based on the ward distance (R package FactoMineR). Groups’ characteristics were compared using the Fisher’s exact test for categorical variables and the Kruskal-Wallis test followed by Dunn’s tests for multiples comparisons for continuous variables. Survival estimates were obtained with the Kaplan-Meier method and statistical tests performed using Cox models. Statistical analysis was performed using R software (www.R-project.org, Vienna, Austria).

## Results

### Comparison of methodologies for determining the clonal architecture of hematopoiesis in a cohort of MPN patients carrying a *JAK2*-V617F mutation

From our cohort of 225 MPN patients positive for the *JAK2*-V617F mutation, we selected 50 patients that carried at least one additional somatic gene mutations by targeted NGS (Figure 1). In these 50 patients we characterized the clonal architecture by cell culture assays, i.e. either single cell liquid cultures or colony assays in methylcellulose, and in 22 of these 50 patients, we also performed single-cell DNA sequencing that allowed to compare the two methodologies (Figure 2A). A single vial of frozen PBMCs was split and unfractionated PBMCs were used for Tapestri scDNAseq, while FACS sorted CD34-positive cells were deposited as single cells into liquid cultures. After 14 days DNA from colonies was extracted and sequenced. scDNAseq of PBMCs allowed scoring a median of 4’367 (range 459-8’009) individual cells per patient sample. The somatic mutations previously defined by bulk NGS were genotyped by scDNAseq in a median of 2’786 cells per sample (range 277-6’717). The lower values mostly corresponded to mutations located in GC-rich regions (e.g. *SRSF2-P95)*. Liquid cultures were performed with CD34-positive cells and a median of 144 colonies per patient (range 27-256) were obtained for analysis. The colony-forming efficiency was 60% (range 14-86%). The VAF of *JAK2*-V617F determined by scDNAseq of PBMCs correlated very well with the VAF measured by allelespecific digital PCR in bulk PBMCs (Figure 2B), whereas VAF derived by genotyping of single colonies showed more variability and an overall trend towards lower values compared to VAF determined in bulk CD34+ cells by allele-specific digital PCR. We suspect that this is due to differences in the efficiencies of colony growth, which may also depend on the presence or absence of additional mutations.

**Figure 1:**
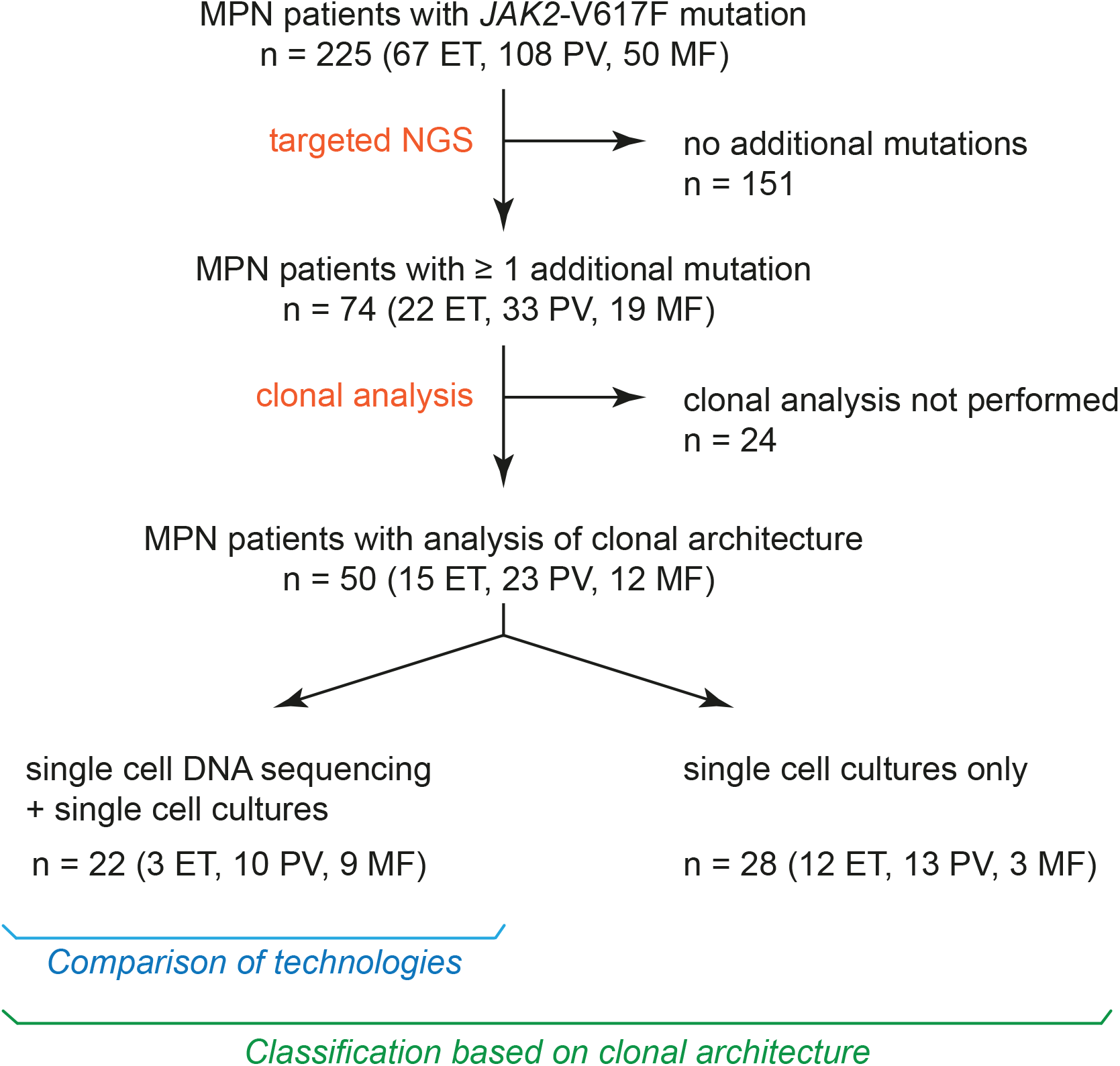
Overview of MPN patients included in the study. A flow chart of patients with myeloproliferative neoplasms (MPN) included in this study and analyses that were performed are shown. NGS, targeted next generation sequencing with a panel of 68 genes. ET, essential thrombocythemia; PV, polycythemia vera; MF, myelofibrosis.

**Figure 2:**
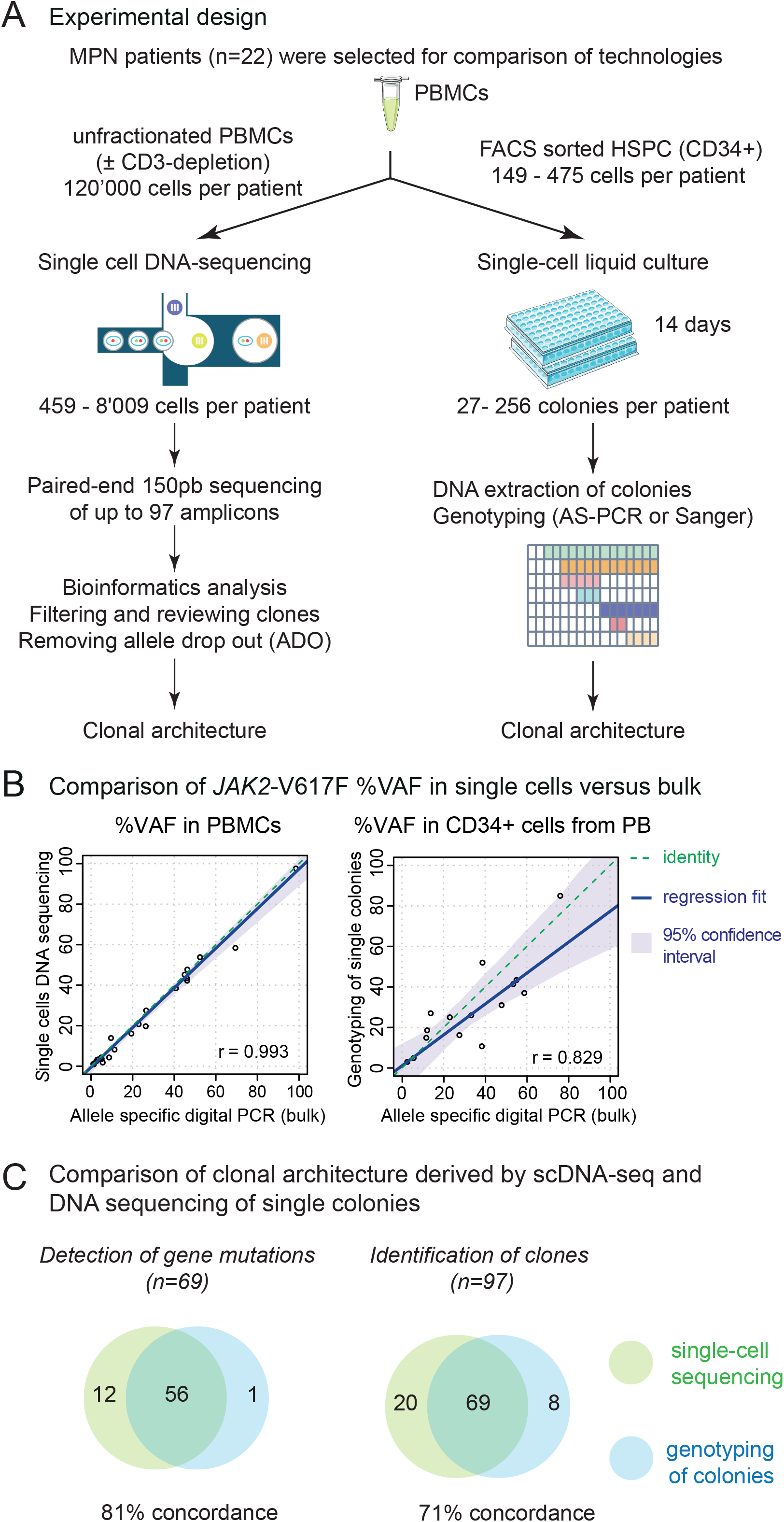
Comparison of methodologies to define the clonal architecture of hematopoiesis in MPN patients carrying more than one somatic gene mutation. (A) Experimental design. Starting material were frozen peripheral blood mononuclear cells (PBMCs). HSPC, hematopoietic stem and progenitor cells; AS-PCR, allele-specific PCR. (B) Correlation between the variant allele frequencies (VAF) measured by digital AS-PCR in bulk cell populations (PBMC or CD34-positive cells), versus VAF deduced from single cell DNA sequencing (scDNAseq) (left panel) or from genotyping DNA from individual colonies (right panel). Non-parametric Passing-Bablok regression was performed and the correlation was estimated by Pearson r parameter. (C) Venn plots showing the concordance and differences in detecting somatic mutations (left panel), or identifying individual clones (right panel) using scDNAseq versus genotyping DNA from individual colonies.

A total of 69 somatic mutations (including *JAK2-*V617F) and 97 clones or subclones were detected by at least one of the two methods (Supplemental Tables S1 and S2). A global concordance of 71% between the two methods was achieved for the detection of individual clones (Figure 2C). Despite the good concordance, some clones were missed by one or the other methodology: 20 subclones in 11 patients were detected by scDNAseq only, and 8 subclones in 5 patients were detected by genotyping of colonies only. In 19/20 cases, the subclones detected only by scDNAseq were due to mutations in 5 different genes not previously scored as mutated by targeted bulk-NGS of granulocyte DNA (Figure 3A and Supplemental Figure S1A). These new gene mutations were found because scDNAseq not only re-sequenced the gene regions where the previously defined mutations lie, but we also amplified and sequenced additional regions, where classical mutational hotspots are located. After reviewing the original NGS alignment files, we found that these newly detected mutations were already present in the granulocyte DNA used for NGS, but with a low VAF (data not shown). Only one subclone with a previously known *EZH2* gene mutation in patient P499A was missed by genotyping of a total of 78 colonies (Supplemental Figure S1C). Thus, genotyping of colonies is very reliable, provided that sufficient number of colonies can be scored, but detection is limited to previously known gene mutations. Tapestri scDNAseq can detect additional subclones because a large number of amplicons (in our case 97) are sequenced in all patient samples.

**Figure 3:**
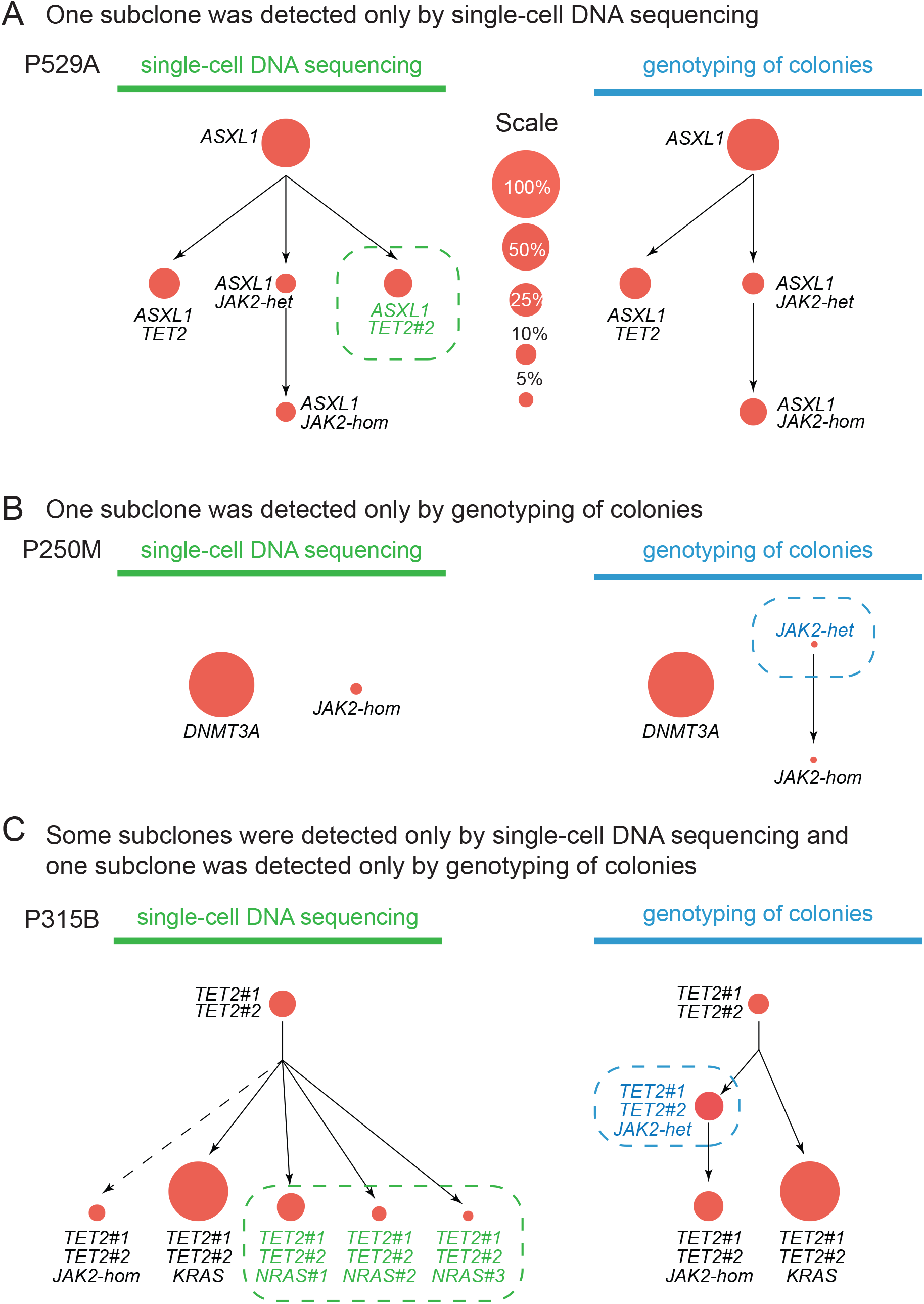
Examples illustrating concordance and discordance in detecting subclones by single-cell DNA sequencing versus genotyping of colonies. Examples of three patients in whom differences in the derived clonal architecture were noted when comparing the two methodologies. The deduced order of acquisition of mutations is drawn from top to bottom. Each subclone is represented by a circle and its size represents the percentage related to the sum of all mutated subclones, which was set to 100%. (A) In P529A one subclone was detected only by single-cell DNA sequencing (marked by dashed green circle). (B) In P250M one subclone was detected only by genotyping of colonies (marked by dashed blue circle). (C) In P315B some subclones were detected only by single-cell DNA sequencing (marked by dashed green circle) and one subclone was detected only by genotyping of colonies (marked by dashed blue circle).

Conversely, genotyping of colonies detected 8 subclones in 5 patients that were not found by scDNAseq. One of these subclones was missed by scDNAseq for technical reasons (insufficient coverage by scDNAseq of the relevant gene region), whereas the other 7 subclones detected only by genotyping of colonies represented intermediate ancestral subclones in the transition from heterozygosity to homozygosity (Figure 3B and Supplemental Figure S1B). Thus, scDNAseq appears to be less reliable at distinguishing heterozygosity to homozygosity. This could be due to an allele drop-out artifact in heterozygous cells. We also used unfractionated PBMCs for scDNAseq, whereas in the single cell liquid cultures the progeny of CD34+ progenitors was scored.

The difficulties distinguishing heterozygosity from homozygosity is inherent to the scDNAseq methodology. To correctly identify cells heterozygous for a mutation, scDNAseq must amplify and sequence the relevant locus from both chromosomes (Figure 4A). If only one of the loci is amplified from a heterozygous cell, then the genotype will be wrongly assigned to either wildtype or homozygous. Allele drop-out typically results in a reduced frequency of reads for the homozygous or wildtype false-positive genotypes (Figure 4B). Identification of allele dropout artefacts was facilitated by detection of the two counterparts of the allele bias resulting in false positive homozygous and wildtype cells occurring at a low but similar frequency (Supplemental Figure S2).

**Figure 4:**
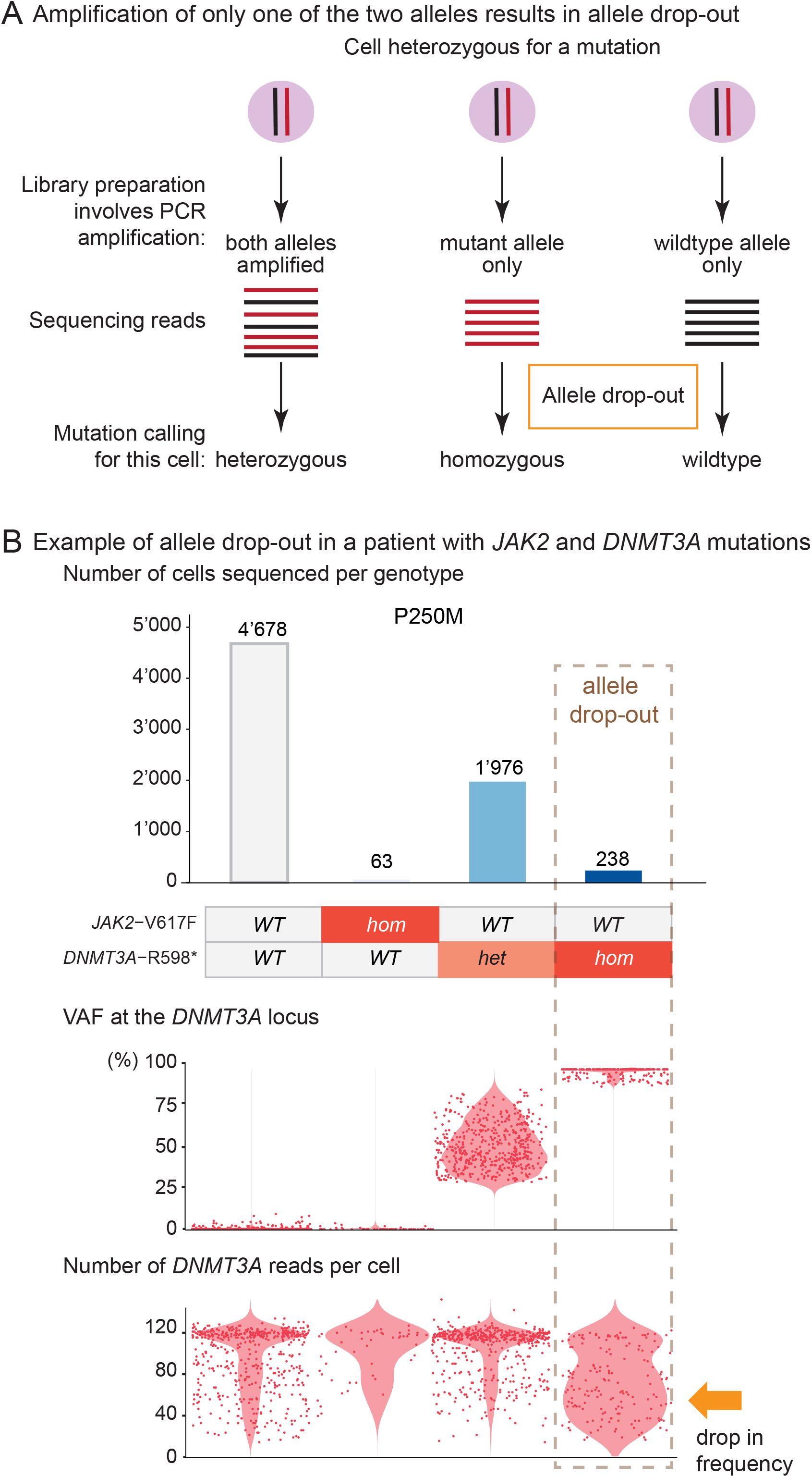
Failure to amplify one of the two alleles by in scDNAseq results in allele dropout artefacts. (A) Allele drop-out is an inherent artefact due to the scDNAseq technology producing from a heterozygous mutation false positive clone with either wildtype or homozygous mutation. (B) Curating the sequencing data allowed to identify and exclude the clones due to allele drop-out. In patient P250M, allele drop-out of a *DNMT3A* heterozygous mutation produced an artifactual homozygous clone which was identify because of a drop in read depth.

To estimate the effects of clonal selection bias caused by the single cell cultures, we performed a detailed analysis in one patient from whom bone marrow cells were available and were able to obtain sufficient numbers of CD34+ to compare the two methodologies using the same starting cell population (Supplemental Figure S3). Heterozygous *JAK2*-V617F mutation was found in 3.1% of CD34+ bone marrow cells by scDNAseq, whereas 10% of colonies (10/102) were heterozygous for *JAK2*-V617F by genotyping of single cell derived colonies. The VAF for *JAK2*-V617F was 5% in peripheral blood granulocytes and 6% in total bone marrow cells. Thus, the higher estimate of *JAK2*-V617F-positive CD34+ progenitors in bone marrow is likely due to increased cloning efficiency of *JAK2*-mutant progenitors compared with wildtype progenitors. Interestingly, the *SF3B1* subclone was slightly underrepresented by genotyping of colonies (15% versus 29%), suggesting that the *SF3B1* mutation decreased the cloning efficiency compared to wildtype progenitors. The presence of *JAK2*-V617F/*SF3B1* double mutant cells/colonies suggests that *JAK2*-V617F was acquired twice in this patient (P368H). A similar pattern was also found in P099D (Supplemental Figure S1).

Overall, our data demonstrate that the two methodologies in most cases produced concordant results yielding the same clonal architectures. Genotyping of colonies may shift the frequencies of subclones when gene mutations increase or decrease cloning efficiencies, whereas scDNAseq can detect small subclones and also additional mutations, but has more difficulties distinguishing heterozygous mutations due to allele drop-out artifacts.

### Correlation between clonal architecture and clinical features of MPN patients with *JAK2-V617F*

To obtain a cohort with sufficient numbers of patients with fully characterized clonal architecture, we combined data from the 22 patients studied by both scDNAseq and genotyping of colonies with data from 28 additional patients studied solely by genotyping of colonies (Figure 1).^9,11^ With these 50 patients, unsupervised clustering using only data on clonal architecture, including 10 parameters listed in Supplemental Methods, defined four well-separated clusters (Figure 5A).

**Figure 5:**
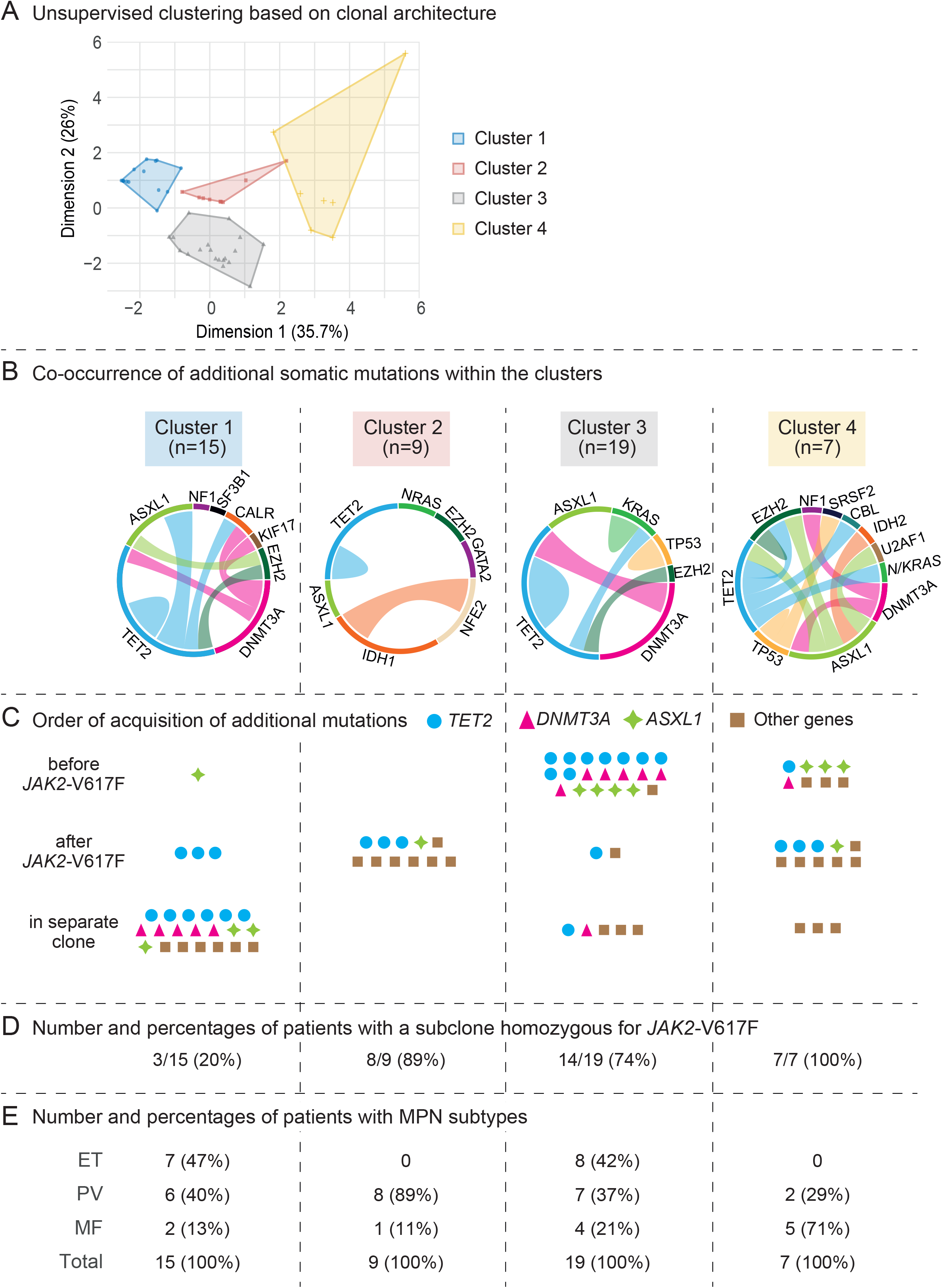
Analysis of clonal architecture allowed defining clusters with specific molecular landscapes. (A) Factor map obtained from the factor analysis with mixed data of clonal architecture parameters (FAMD analysis). The 4 clusters obtained after a clustering based on the ward distance are indicated by colored areas. (B) Circos plots illustrating co-occurrence of somatic mutations in the same individual. The length of the arc corresponds to the frequency of the mutation, whereas the width of the ribbon corresponds to the relative frequency of cooccurrence of 2 mutations in the same patient. (C) Order of acquisition of additional somatic mutations shown in relation to the *JAK2-*V617F driver mutation for each of the 4 clusters. Each symbol represents one gene mutation, as indicated at the bottom. (D) Proportion of patients with at least one subclone with homozygous *JAK2*-V617F mutation detectable in each cluster. (E) Proportions of MPN subtypes across clusters.

The clusters did not differ in the distribution of the individual mutated genes, but cluster 2 showed the least complex mutational landscape and cluster 4 had the most complex profile with the largest number of additional mutations per patient (Figure 5B). The order in which the additional mutations were acquired with respect to JAK2-V617F also differed between clusters. In cluster 2, *JAK2-*V617F was always the first event, whereas in cluster 3 at least one additional mutation always preceded the acquisition of *JAK2*-V617F (Figure 5C). In cluster 1, the majority of additional mutations represented a clone separate from *JAK2-*V617F. Cluster 4 was more complex, without a strict order of events. The clusters also differed in the presence or absence of subclones homozygous for *JAK2-*V617F. In cluster 1, only 20% of patients had a homozygous subclone, whereas in cluster 4 a homozygous subclone was detectable in all patients (Figure 5D).

The patients assigned to the four clusters also showed differences in clinical features. Patients with ET were present only in clusters 1 and 3 (Figure 5E), whereas patients with PV and MF were found in all four clusters, but MF was most frequent in cluster 4 (Figure 6A). Patients assigned to cluster 3 were on average older at the time of diagnosis than patients in clusters 1 or 2 (Figure 6B). Patients in cluster 4 had a reduced overall survival (OS) with a median OS of 5 years (Figure 6C). The hazard ratio (HR) for death in cluster 4 *vs* the other clusters was 5.32. Multivariate analysis confirmed this result, and cluster 4 was associated with reduced overall survival independent of the subtype of MPN diagnosis and independent of the age at the time of diagnosis (Supplemental Table S3). Evolution to AML or myelodysplastic syndrome was more frequently observed in patients assigned to cluster 4 (29% of patients) than in patients from other clusters (7%, 11% and 11% for clusters 1, 2 and 3, respectively). Interestingly, few deaths occurred in patients of clusters 1 and 2 despite the presence of high-risk additional mutations, suggesting that such mutations occurring in a separated clone or isolated in the driver clone may have a distinct prognostic impact. The subtype of MPN and the total number of additional mutations also had a prognostic impact in this cohort (Figure 6D). However, we found that clonal architecture classification performed better for predicting OS as demonstrated by a higher C-index (0.72 *vs* 0.68 and 0.67, respectively).

**Figure 6:**
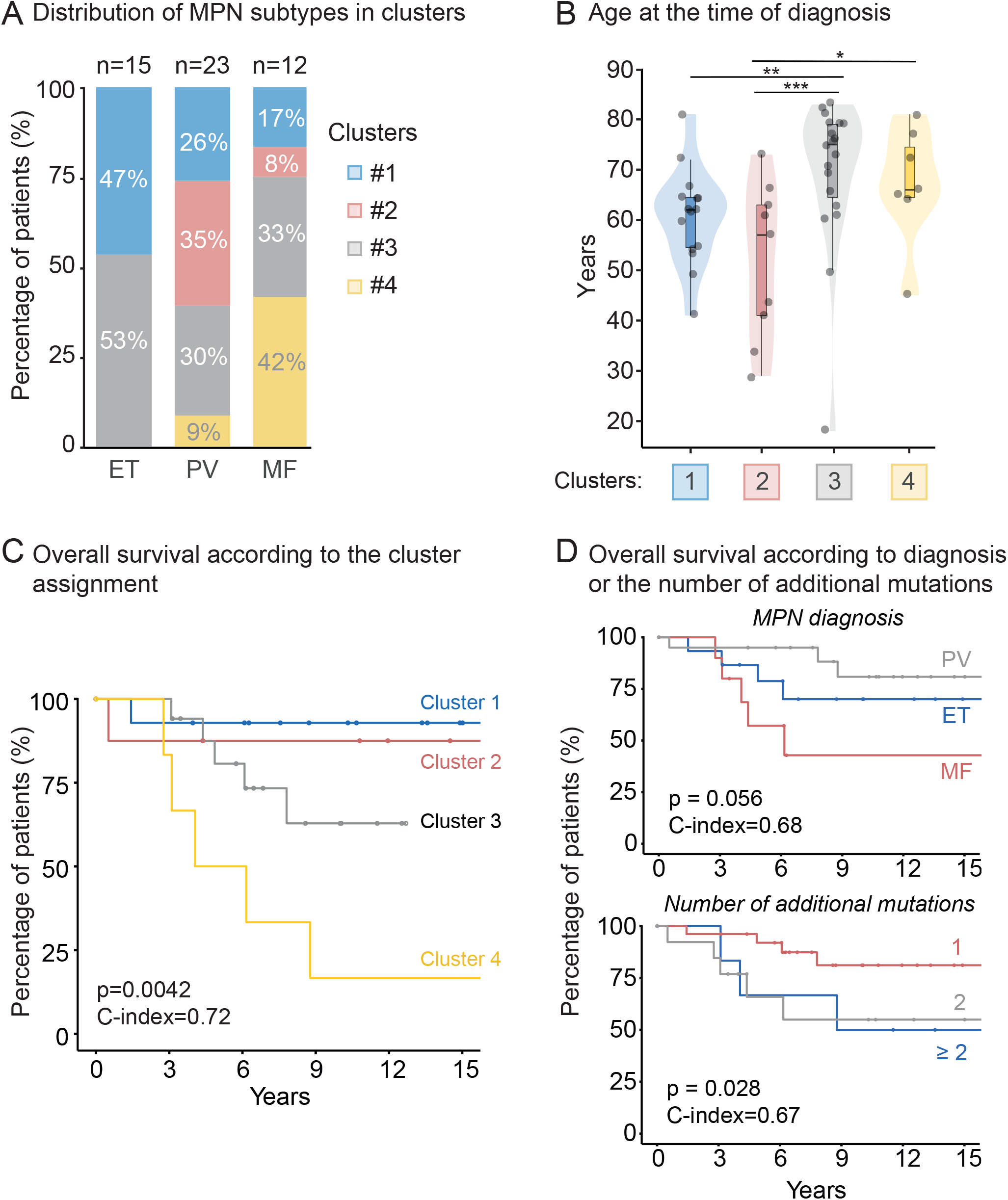
Clinical features of MPN patients assigned to clusters derived by clonal architecture data. (A) Frequencies of the four clusters within *JAK2*-V617F-positive MPN patients diagnosed as ET, PV or MF. (B) Age distribution at the time of diagnosis. Differences were tested by a Kruskal-Wallis test followed by Dunn’s multiples tests. (C). Overall survival of MPN patients assigned to the four clusters. D) Overall survival of MPN patients according to MPN diagnosis (top), and number of additional gene mutations (bottom). Survival estimates were obtained with the Kaplan-Meier method and statistical tests performed using a logrank test. The performance of prediction was accessed by Harrell’s concordance index (C-index).

## Discussion

The clonal architectures deduced from genotyping of colonies and from Tapestri scDNAseq of PBMCs showed good overall concordance (Figure 2C). Genotyping by scDNAseq showed higher sensitivity for mutations with low VAF due to the high number of single cells that were scored per patient sample. Furthermore, scDNAseq also allowed identifying subclones defined by additional mutations not previously found by bulk NGS, because the methodology allows to include other genes or gene regions (e.g. mutational hot spots) covered by sequencing and not only the location where the previously identified mutation lies (Figure 3). As expected, scDNAseq had more difficulties to distinguish between heterozygous and homozygous mutations (Figure 4). Allele drop-out artefacts are inherent to the scDNAseq methodology and their rate varies for different gene loci and between samples, in most cases ranging around 3 to 10% of individual cells.^14^ By a manual curation, we identified and removed clones corresponding to allele drop-out, but we cannot exclude that some cells in these clusters could be true homozygotes. Another reason why some of the results between scDNAseq and genotyping of colonies differed could be that different starting cell populations were used for the two methodologies, i.e. unfractionated PBMC versus peripheral blood CD34+ progenitors.

Other approaches that do not require expansion of cells in culture and can be applied to unfractionated cell populations have been developed for genotyping. The 10x chromium method uses RNA from single cells as template for sequencing the mutated part of the gene. The sensitivity is dependent on the expression levels of the corresponding mRNA and decreases with the distance from the 3’-end of the mRNA. The sensitivity can be improved by adding nested PCR for amplifying cDNA of the specific locus. This “GoT” approach applied to MPN allowed genotyping for *CALR* mutations, but did not work reliably for *JAK2*-V617F mRNA,^15,16^ which is expressed at lower levels than *CALR* and located far from the 3’-end.

Several methods for genotyping of FACS-sorted single cells have been described that are less suited for high throughput applications, because each cell has to be processed individually. A DNA based approach that involved genome-wide DNA amplification of single cell DNA followed by Taqman based detection of previously defined mutations was applied to PMF patients.^17^ The Smart-seq2 method used full-transcript RNA sequencing that avoids bias for the 3’-ends, but produced high rates of allele drop-out due to the non-random monoallelic nature of expression of the selected genes in single cells.^18^ TargetSeq, a protocol combining fulltranscript RNA sequencing with DNA targeted genotyping in individual FACS-sorted cells resulted in a good efficiency of genotyping and low rate of allele drop-out,^19^ but the number of individual cells and transcriptomes that can be analyzed is lower than with the GoT. The Tapestri scDNAseq methodology using encapsulation allowed genotyping several thousand single cells per patient sample with an acceptable rate of allele drop-out and is suited for high throughput applications. However, this DNA-based methodology does not allow simultaneous sequencing of the transcriptome. Currently, the high costs of this methodology is a limiting factor for large-scale use.

These technologies now allow to more efficiently determine the clonal structure of hematopoiesis in patients with multiple somatic mutations and allow to address to what degree the clonal architecture influences clinical features and outcome. We defined 4 distinct clusters by unsupervised analysis of data on clonal architecture from 50 MPN patients that were genotyped by Tapestri scDNAseq and/or single cell colony assays. These four clusters differed from each other by the order of acquisition of the mutations, the total number of mutations, the complexity in branching and evolution of the subclones and the presence or absence of a subclone homozygous for *JAK2*-V617F (Figure 5). All ET patients were found in either cluster 1 or 3. In cluster 1, the majority of additional mutations represented a clone separate from *JAK2-*V617F, whereas in cluster 3 at least one additional mutation always preceded the acquisition of *JAK2-V617F* (Figure 5C).

Previous reports focusing on *TET2* or *DNMT3A* also found that in the majority of ET patients the *TET2* or *DNMT3A* mutation preceded the acquisition of *JAK2*-V617F, but in these reports no bi-clonal patterns in ET were described.^12,20^ In cluster 2, *JAK2*-V617F was always the first event, and these patients were on average younger than those assigned to the other clusters (Figure 6B). Patients assigned to cluster 4 had always a detectable subclone homozygous for *JAK2*-V617F and overall carried a larger number of additional mutations without a preferred order of acquisition resulting in more complex subclonal structures than the other clusters.

A global pan-MPN genomic classification for prognosis stratification was proposed.^21,22^ Our clonal architecture-based cluster 4 was a risk factor for predicting reduced overall survival and in multivariate analysis was independent of the MPN subtype and the age at diagnosis. The number of additional mutations was previously associated with worse prognosis in patients with MPN in general,^9,23^ and PMF in particular,^9,23^ but in our study the cluster classification performed better in predicting overall survival than the number of additional mutations (Figure 6D). High molecular risk (HMR) mutations with strong predictive power in PMF, i.e. mutations in the genes *ASXL1, EZH2, IDH1/2, SRSF2, or U2AF1*,^10,22,24^ were present in all 4 clusters and were not exclusive for cluster 4. Our results suggest that mutations in HMR genes occurring in a separated clone (here mostly in cluster 1) may not be associated with an adverse prognosis. The cooperative effects of additional mutations with *JAK2*-V617F has been documented in several mouse models, but depended on being present in the same hematopoietic cell.^25–29^

Our results suggest that deciphering the clonal architecture in patients with MPN that carry multiple gene mutations can improve the molecular prognostic stratification that until now was primarily based on the number and type of gene mutations.

## Acknowledgements

We thank Stella Stefanova, Lorenzo Raeli and Emmanuel Traunecker from the flow cytometry core facility of Department of Biomedicine for help in cell sorting, Christian Beisel from the Genomics Facility Basel (ETH Zurich, D-BSSE located in Basel operated jointly with the University of Basel.) for sequencing support, and members of the laboratory for helpful discussions and critical reading of our manuscript.

