## Supplemental Figures for "Impact of clonal architecture on clinical course and prognosis in patients with myeloproliferative neoplasms"

### Supplemental Figure S1 (related to Figure 3)

#### A Patients with subclones detected only by single-cell DNA sequencing of PBMCs and not previously detected by targeted NGS

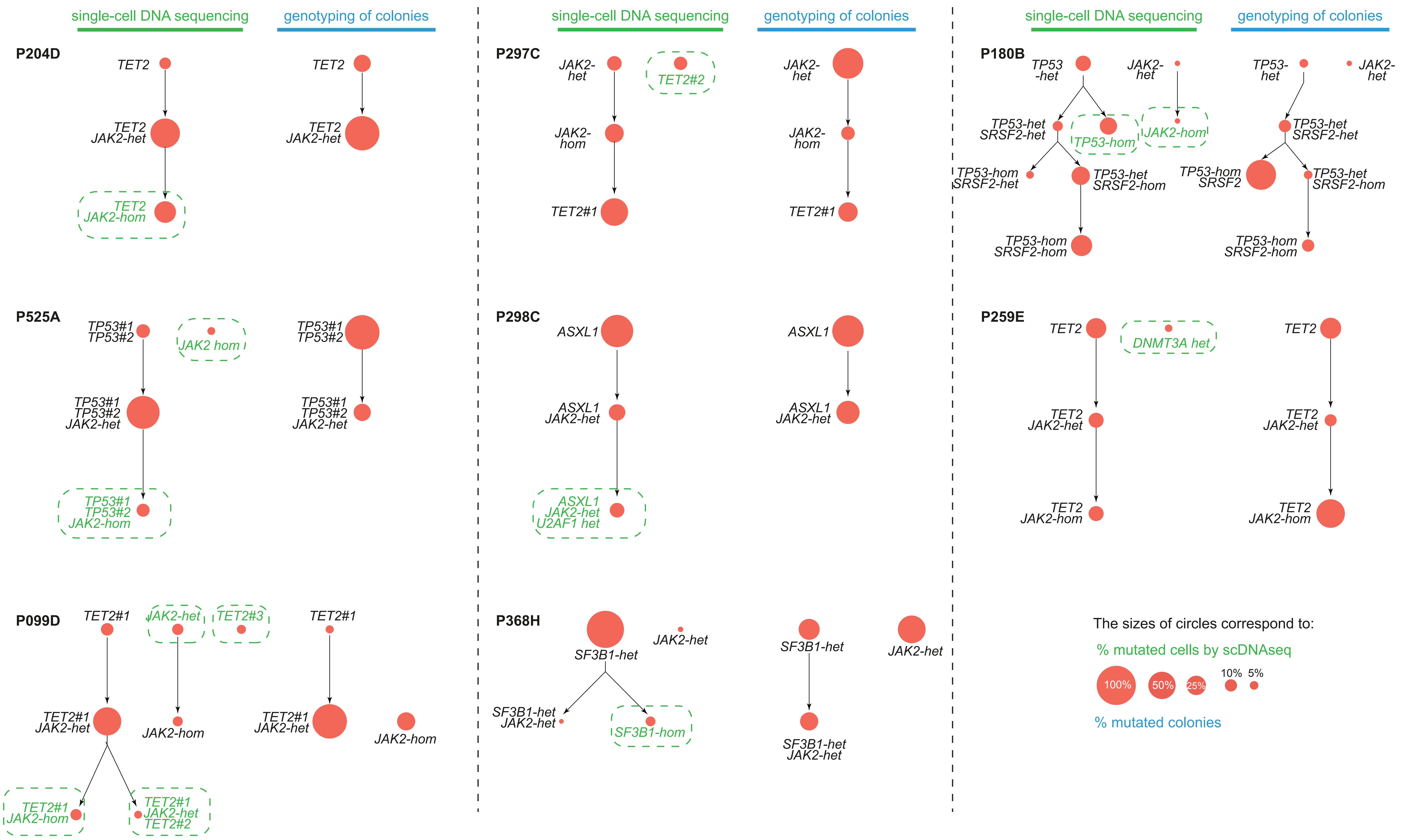


#### B Patients with subclones detected only by genotyping of colonies

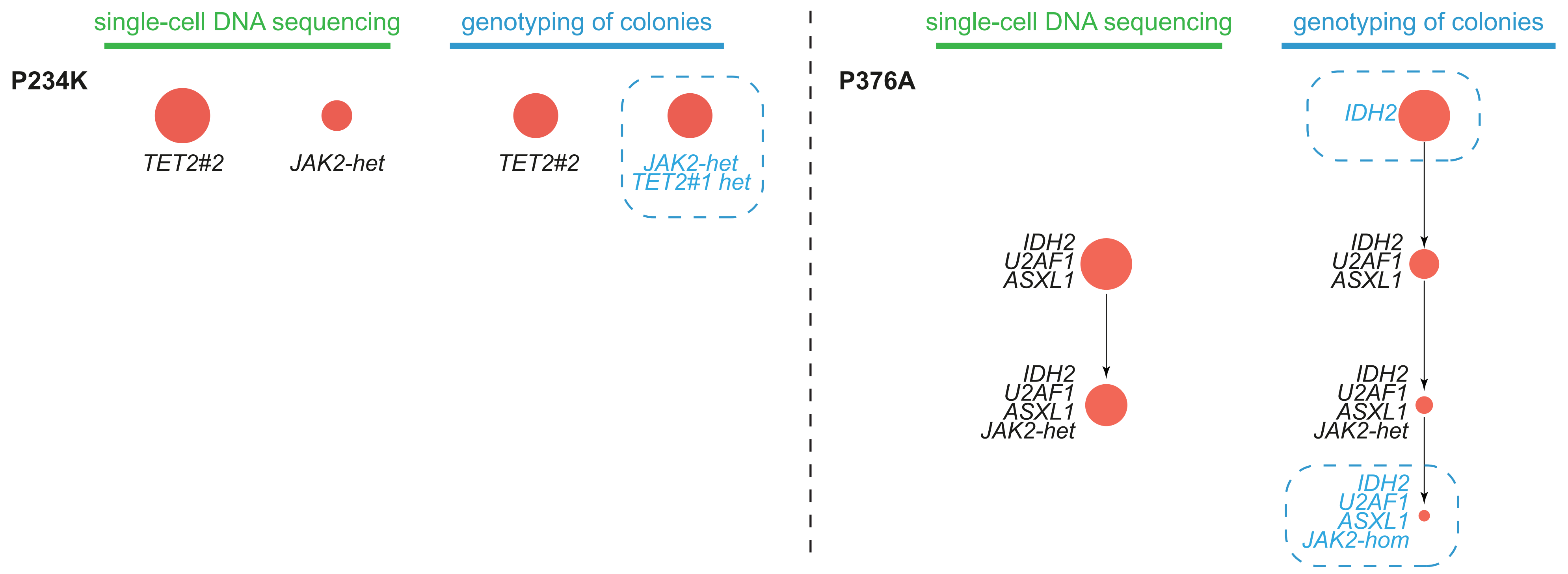


#### C Complex case with some subclones detected only by single-cell DNA sequencing and others detected only be genotyping of colonies

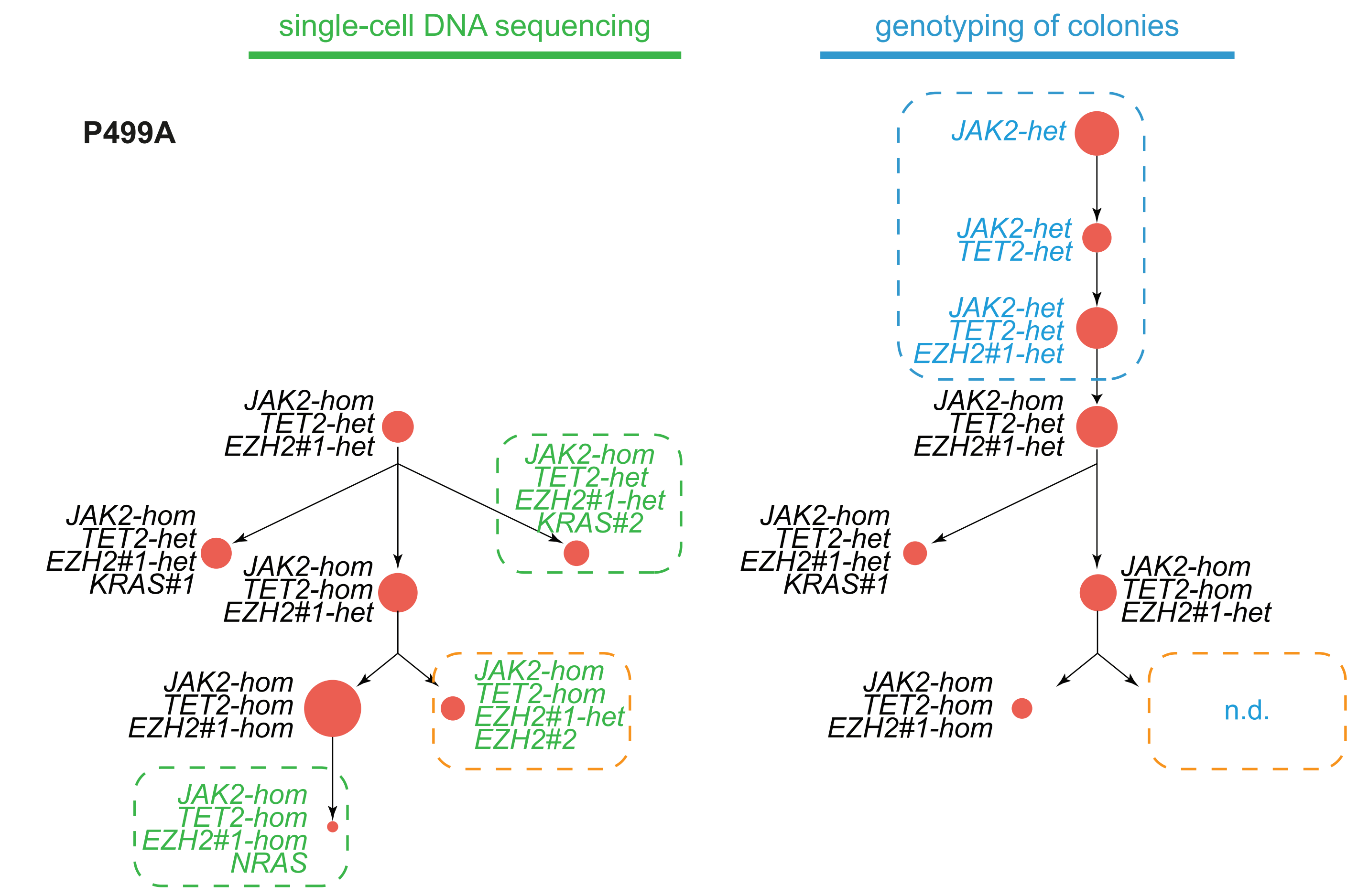


**Legend to Supplemental Figure S1: Comparison of clonal architecture derived from from scDNAseq of PBMCs and genotyping of colonies.** The order of acquisition of mutations is drawn from top to bottom. Each subclone is represented by a circle and its size corresponds to the percentage related to the sum of all mutated subclones, which was set to 100%. Clones highlighted in green or blue are clones detected only by one of the two methods (green for scDNA-seq and blue for genotyping of colonies). Note that all subclones found only by scDNAseq and shown in (A) represent new mutations not previously detected by bulk NGS, and only one subclone with a previously known *EZH2* gene mutation in patient P499A was missed by genotyping of colonies in (C).

### Supplemental Figure S2 (related to Figure 3)

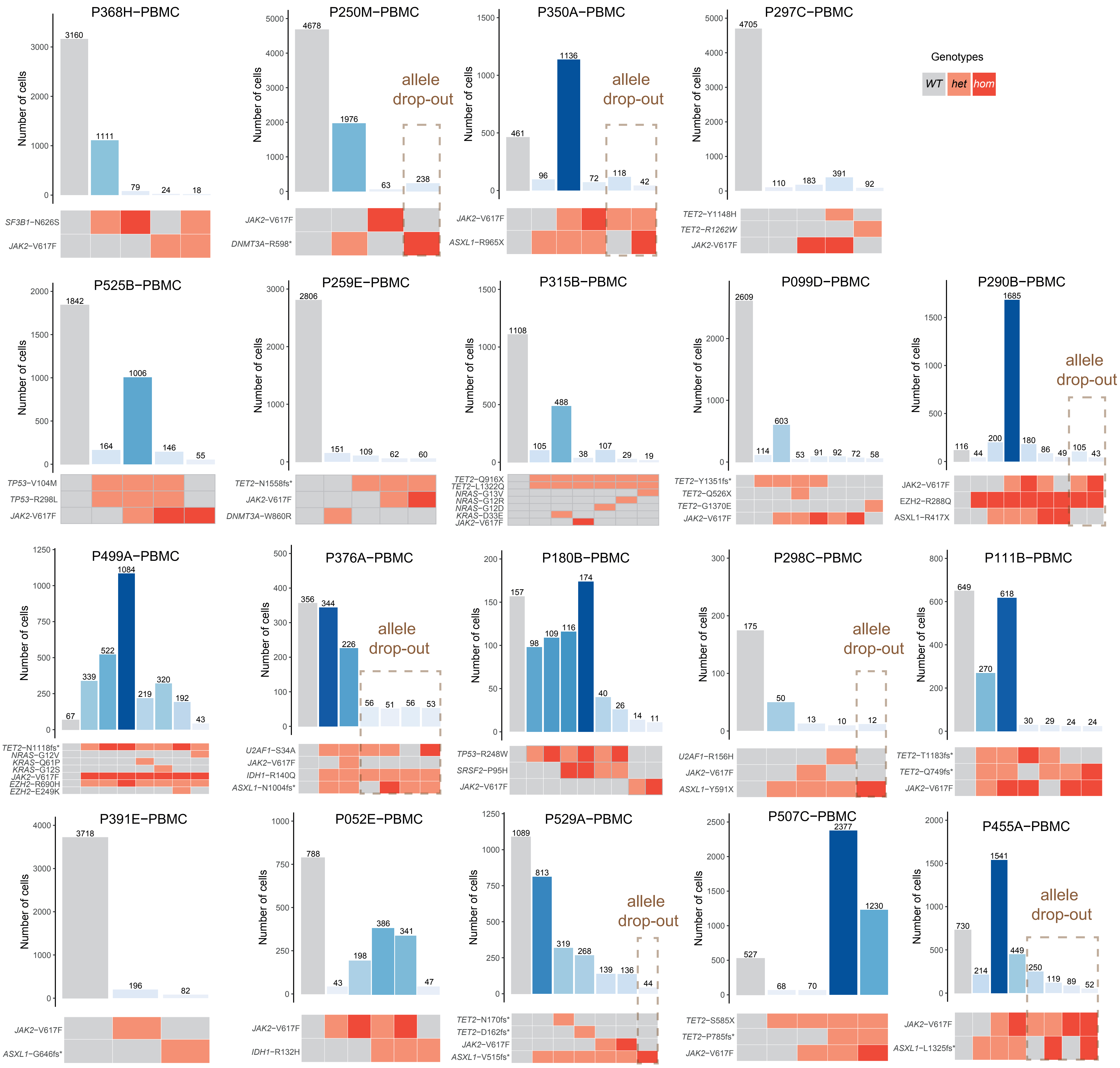


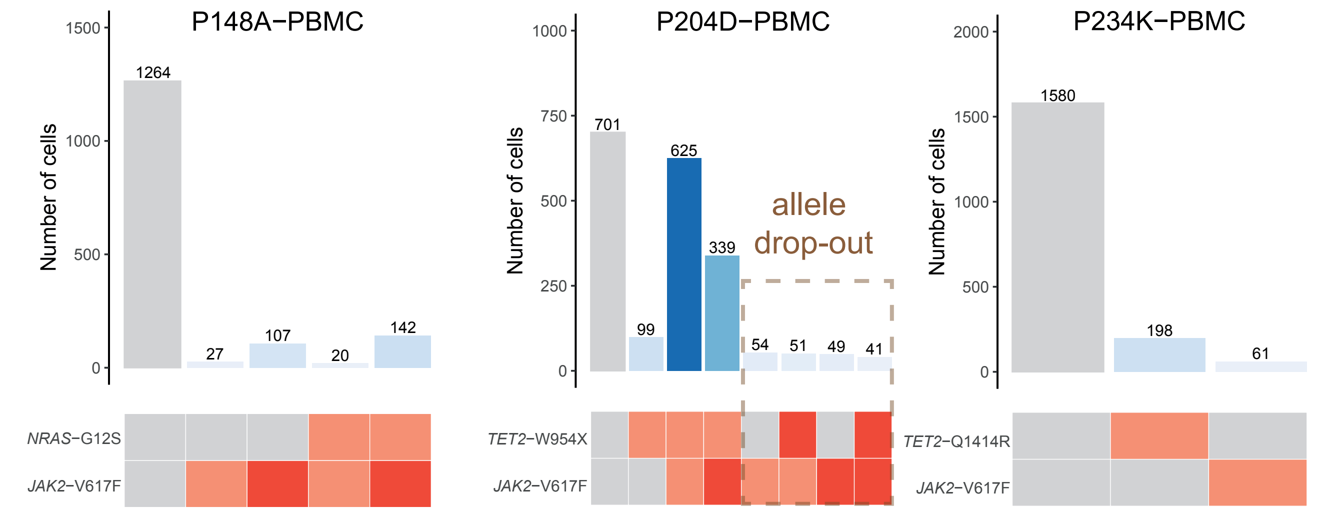


**Legend to Supplemental Figure S2:** Clonal architecture analysis of selected MPN samples using scDNA-seq. Each sample is presented with the number of cells assigned to a separated clone (top plot). Genotypes of clones are represented by the heat map (bottom panel), where mutations are colored according to their zygosity. Artefactual clones identified as allele drop out are represented by a grey square.

**Supplemental Figure S3 (related to Figure 3)**

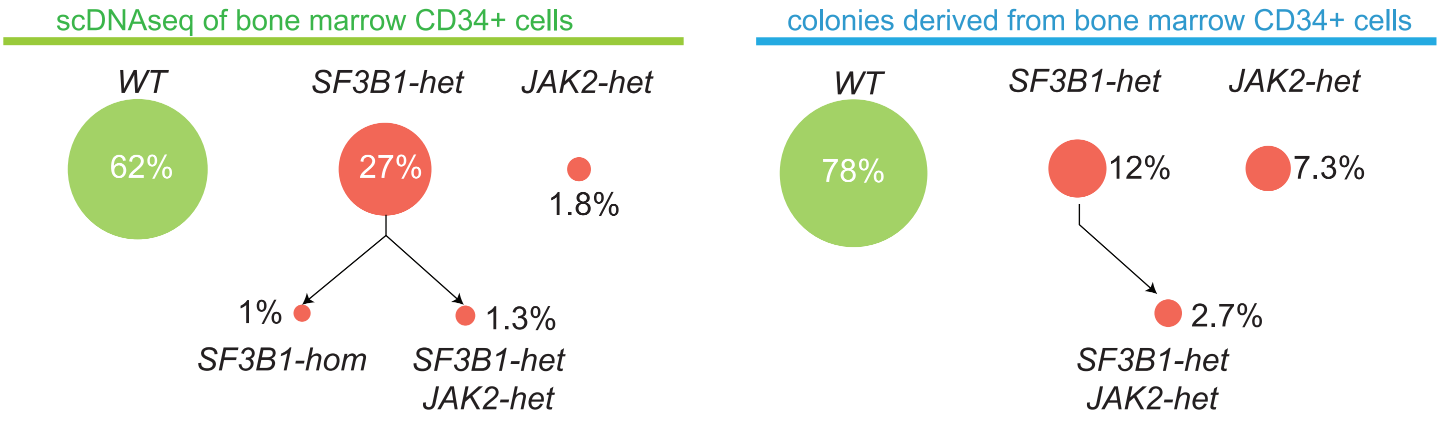


**Legend to Supplemental Figure S3.** Comparison of scDNA-seq and genotyping of colonies for CD34+ bone marrow cells of patient P368H. Both single-cell DNA sequencing and single-cell liquid culture followed by genotyping of colonies were performed on FACS-sorted CD34-positive cells from a bone marrow sample. Each circle represents the percentage related to the sum of all cells or colonies genotyped. Wildtype cells were represented by a green circle and mutated clones by red circles.
